# In situ identification of secondary structures in unpurified *Bombyx mori* silk fibrils using polarized two-dimensional infrared spectroscopy

**DOI:** 10.1101/2022.05.11.491460

**Authors:** Giulia Giubertoni, Federico Caporaletti, Steven Roeters, Adam S. Chatterley, Tobias Weidner, Peter Laity, Chris Holland, Sander Woutersen

## Abstract

The mechanical properties of biomaterials are dictated by the interactions and conformations of their building blocks, typically proteins. Although the macroscopic behaviour of biomaterials is widely studied, our understanding of the underlying molecular properties is generally limited. Among the non-invasive and label-free methods to investigate molecular structures, infrared spectroscopy is one of the most commonly used tools, because the absorption bands of the amide groups strongly depend on protein secondary structure. However, spectral congestion usually complicates the analysis of the amide spectrum. Here, we apply polarized two-dimensional (2D) infrared spectroscopy (IR) to directly identify the protein secondary structures in native silk filks cast from *Bombyx mori* silk feedstock. Without any additional analysis, such as peak fitting, we find that the initial effect of hydration is an increase of the random-coil content at the expense of the *α*-helix content, while the *β*-sheet content is unchanged, and only increases at a later stage. This paper demonstrates that 2D-IR can be a valuable tool for characterizing biomaterials.

## Introduction

The mechanical properties of multi-scale hierarchical biomaterials, such as the rigidity of bones or the toughness of spider silk, are dictated by the molecular properties of their building blocks that self-assemble to form ordered hierarchically supramolecular structures. ^1,2^ Among the different proteins that act as biomolecular building blocks, silk proteins (i.e. fibroin from silkworms or spidroins from spiders) are among the most extensively studied and have become a model biopolymer system, as a result of their accessibility for research and outstanding mechanical and biocompatible properties. ^3^ Silk proteins self-associate to form hetero-nano composite and highly hierarchical supramolecular fibrils. These fibrils are composed of ordered nano-crystals embedded in disordered amorphous regions. ^4^ The molecular properties of the fibrils, such as the adopted secondary structure and the crystal size, ^5^ dictate the mechanical properties of silk. The *β*-sheet appears to be the most stable form and dominates the crystalline content of natural silk fibres, ^6^ while helical and random structures can be observed in the non-crystalline material.^7^ Less stable *α*-helical and random coil structures dominate films cast from water by evaporation under mild conditions, assumed to reflect the prevalence of random coil conformations in aqueous solution.^8–12^ Upon exposure to high humidity, the increased water content causes a glass transition induced softening and an increase in extensibility, as *α*-helices can be converted to swollen random coil structures, while the *β*-sheet content can increase.^7^ Multiple techniques have been applied to study the silk-protein structure under different humidity conditions, providing a clear understanding of the adopted conformation in the nano-crystals. However, the molecular arrangement in the amorphous regions is still debated, principally due to the fact that most structure-sensitive techniques identify these regions only by omission, or samples require harsh treatments that can strongly affect and change the protein structure.

This problem can be avoided by using infrared spectroscopy, which is a label-free and noninvasive technique, widely used to investigate the structure and conformation of biomolecular building blocks.^13^ The molecular vibrations of the amide groups, in particular the amide I mode that involves the carbonyl stretching (Fig. 1a), are sensitive to the protein conformation. In *β*-sheet and *α*-helix structures, amide groups are connected by hydrogen-bonds, leading to a long-range order along the protein backbone and couplings between the amide vibrations, which are mostly of a dipolar nature.^13–17^ These couplings give rise to delocalized normal modes, and for both ideal *β* sheets and *α* helices, the two most important IR-active normal modes have perpendicular transition dipole moments.^13,18^ In the case of anti-parallel *β*-sheets,^13,16,19^ the so-called A_⊥_ and A_‖_ modes absorb at 1620–1630 cm^*−*1^, and 1680–1700 cm^*−*1^, respectively. ^13,16,20^ In the case of an *α*-helix, the frequency difference between the parallel (A) and the perpendicular (E) modes is typically only few wavenumbers,^18,21^ and the *A* and *E* bands have significant overlap (Fig. 1b), resulting in a single band centered at *∼*1650-1660 cm^*−*1^.^13^

**Figure 1:**
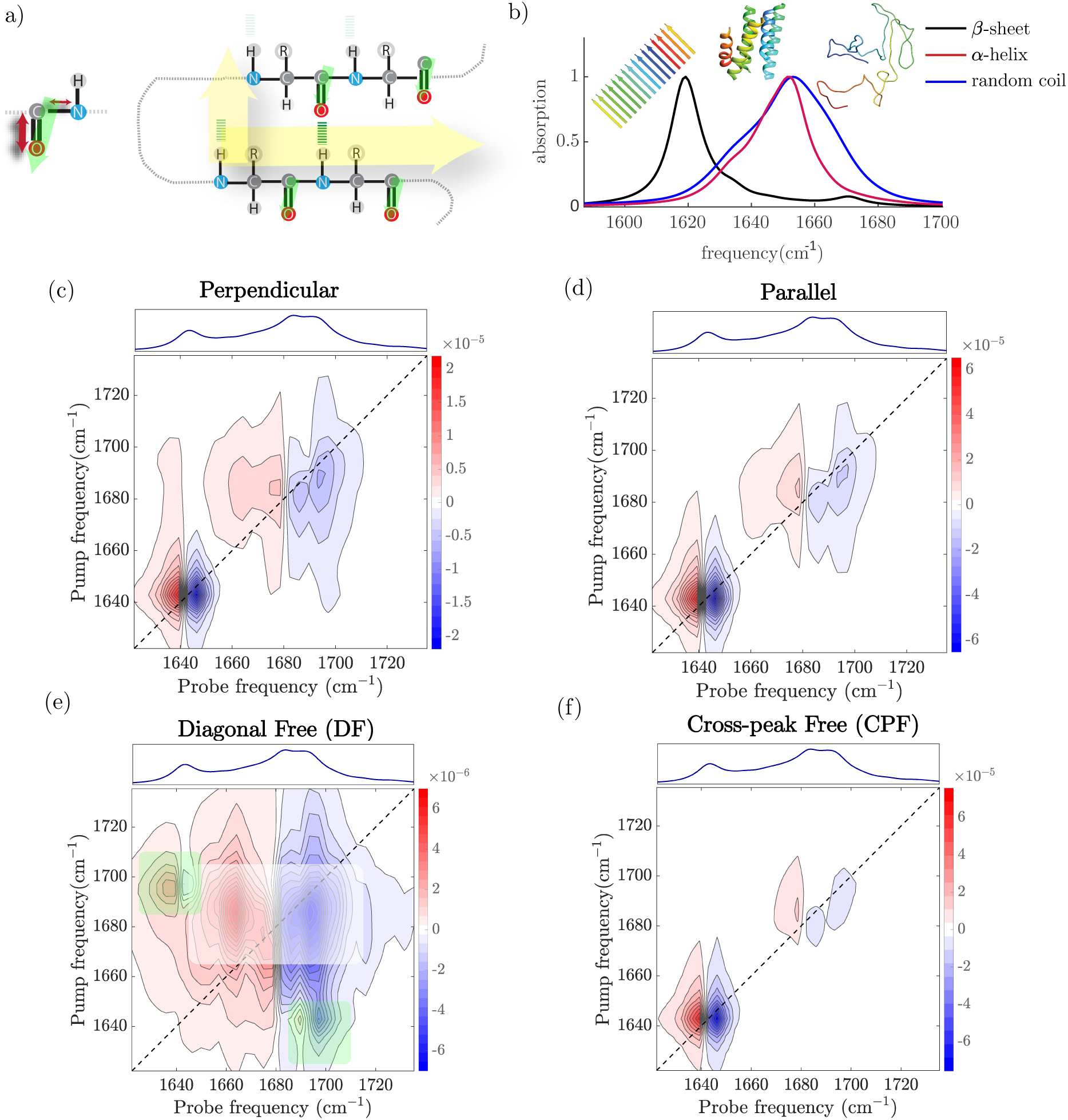
a) Schematic of amide I vibrational modes in a single molecule and in a series of coupled amide groups. b) Simulated linear spectra of short peptides. c)-f) Simulated 2DIR spectra for a a short peptide adopting *β*-sheet and *α*-helix conformation. Color coding: Δ*A <* 0 blue, Δ*A >* 0 red. e) Diagonal free 2DIR spectrum is obtained by subtracting the (3 *×*) scaled perpendicular (c) to parallel (d), while cross-peak free spectrum (f) is obtained by subtracting parallel (d) to two times perpendicular (c). The green and white rectangles in (e) show the position of the cross-peaks associated to the *β*-sheet and *α*-helix secondary structures, respectively.

Unfortunately, the infrared absorption spectra in the amide I region are generally rather congested, limiting our ability to disentangle the underlying amide I bands in an unambiguous manner. Many indirect methods of data analysis have been developed to solve this problem.^22^ However identifying secondary structures based only on the the absorption frequencies obtained via fitting may not be sufficient to determine the secondary structures present. Indeed, an assignment based on the absorption frequency is not always unique because of secondary effects, such as solvent interactions, that may shift the amide I vibrational frequencies and more specific spectral signatures are required. To overcome these problems, we use two-dimensional infrared spectroscopy (2DIR) to investigate the secondary structure of silk. 2DIR spectroscopy can detect couplings between vibrational modes directly, and use these to obtain structural information^23–29^ in a manner somewhat similar to 2D-NMR, where spin-spin couplings are detected and used to obtain structural information. ^30^ 2D-IR Spectroscopy is particularly well suited to study time-dependent changes in the secondary structure of proteins. ^26,29,31,32^

Here, we apply 2DIR spectroscopy to investigate *in situ* the secondary structures present in films produced using unpurified fibroin from *Bombyx mori* silkworms. By selecting specific polarization combinations, we obtain unique spectral signatures that allow us to disentangle and assign vibrational bands to specific secondary structures. We find that the film contains *α*-helical, *β*-sheet and random-coil structures, with *α*-helical structure being most predominant at ambient humidity. The exposure of the film to a saturated water environment leads to a decrease of the *α*-helix contribution and an increase in the spectral region of the *β*-sheet and random coil conformations. This work demonstrates that 2DIR spectroscopy can be used, without peak fitting, to measure structure conversion in a hydrated silk film. Thus, we highlight the unique ability of 2DIR to characterize the protein secondary structure in biomaterials in a direct manner, opening a new set of potential interdisciplinary applications for 2DIR.

## Methods

### Silk-film preparation

Films were prepared using the native silk feedstock (NSF) from the middle-posterior (MP) sections of silk glands from commercially reared *Bombyx m*. silkworms (four-way poly-hybrid cross of two Japanese and two Chinese strains) in their 5th instar. Specifically, silkworms during the early stages of cocoon construction were sacrificed by decapitation, allowing the two silk glands and haemolymph to be ejected into a petri dish.

One gland was selected and transferred to a second petri dish and immersed in type I (distilled and deionised) water. Using a pair of tweezers, the gland was divided around the mid-point and the anterior portion (containing more sericin) was discarded. A second cut was made where the (wider) middle section started and the (relatively narrow) posterior section was also discarded. The thin membrane was peeled off the MP gland section, using fine tweezers under a stereomicroscope, and the viscous NSF (around 0.15 g, containing around 0.035 g of predominantly fibroin) was transferred to a polystyrene weighing boat. Around 2 to 3 mL of type I water was added, the weighing boat was loosely covered with tissue paper and allowed to stand at ambient temperature. The NSF initially dissolved into the water, then a film formed as the water evaporated. The film was allowed to dry under ambient conditions for a few days, before being transferred to a vacuum oven (still in the weighing boat). Drying to constant weight was completed over several hours at 60°C under vacuum. Then the film was peeled off the weighing boat and transferred to a sealed plastic bag for storage until required.

### FTIR and 2DIR Spectroscopy

Samples for infrared spectroscopy were prepared by selecting a *∼*1 cm^2^ part of a *∼*10*µ*m thick silk film, and placing it between two CaF_2_ windows in a customized cylindrical sample holder. After the measurements at ambient humidity (*<* 60%), the sample was placed for 120 minutes in a D_2_O environment at a relative humidity of *∼* 85%. A Perkin-Elmer Spectrum-Two FTIR spectrometer (resolution 4 cm^*−*1^) was used to measure the FTIR spectra. A detailed description of the setup used to measure the 2DIR spectra can be found in ref. Briefly, pulses of wavelength 800 nm and with a 40 femtosecond duration are generated by using a Ti:sapphire oscillator, and further amplified by using a Ti:sapphire regenerative amplifier to obtain 800 nm pulses at 1 kHz repetition rate. These pulses are then converted in an optical parametric amplifier to obtain mid-IR pulses (*∼*20 *µ*J, *∼*6100 nm) that has a spectral full width at half max (FWHM) of 150 cm^*−*1^. The beam is then split into a probe and reference beam (each 5%), and a pump beam (90%) that is aligned through a Fabry-Pérot interferometer. The pump and probe beams are overlapped in the sample in an *∼*250-*µ*m focus. The transmitted spectra of the probe (*T*) and reference (*T*_0_) beams with pump on and off are then recorded after dispersion by an Oriel MS260i spectrograph (Newport, Irvine, CA) onto a 32-pixel mercury cadmium telluride (MCT) array. The probe spectrum is normalized to the reference spectrum to compensate for pulse-to-pulse energy fluctuations. The 2DIR signal is obtained by subtracting the probe absorption in the presence and absence of the pump pulse. Parallel and perpendicular 2DIR spectra are recorded by rotating the pump beam in a 45° angle with respect to the probe beam and selecting the probe beam component that is either perpendicular or parallel to the pump beam using a polarizer after the sample. To minimize pump-scattering contributions, we measured the average of two photoelastic modulator (PEM)-induced pump delays, such that the interference between the scattered pump beam and the probe beam has a 180° phase in one delay with respect to the other delay.

### Spectral calculations

The spectral calculations were performed in accordance with the formalism described in Ref. 25,34. We use the transition-dipole coupling (TDC) model^35^ to determine the couplings between the amide-I modes in the protein backbones. In this model, the through-space coupling is approximated by a Debye-like coupling that is dependent on the relative orientation and distance between the amide-I oscillators. The orientation and magnitude of the isolated amide-I transition-dipole moments have been determined with a DFT (Density Functional Theory) calculation.

First, we apply this formalism to three different structures to calculate ‘pure-component’ 1D- and 2D-IR spectra for three types of secondary structure: random coils obtained from a molecular-dynamics simulation of *α*-synuclein,^36^ from X-ray crystallography experiments on the mainly *α*-helical structure of spindroid proteins, a component of from spider silk, (PDB: 3LR2^37^) and from a perfect antiparallel *β*-sheet structure generated in Chimera. ^38^ From the 73 *µ*s *α*-synuclein simulation, we took 73 snapshots (frames), starting from frame 1, spaced by 1 *µ*s of simulation time (see Fig. 1B for the first frame). We then calculated the 2DIR spectra for each of the frames, and determined the average (2D)IR spectrum of the ensemble, in order to optimally sample the many conformations that a randomly-coiled structure adopts. The *α*-helical (2D)IR spectra were calculated by taking just the *α*-helical structure of chain A of PDB 3LR2 and removing the three prolines from the sequence, together with the 2-3 residues after the proline residue at the ends of the three proline-containing helices. This latter step was performed to keep the ‘pure component’ calculations as clean as possible; the amide-I local mode of proline is red-shifted by *∼*19 wavenumbers,^34^ which leads to more complex amide-I normal modes. Finally, the perfect antiparallel *β*-sheet was created by placing 16 poly-alanine 16-mers in an antiparallel sheet with a 5 Å spacing, by rotating every second monomer by 180 degrees and shifting it by 1.5 Å to align the amide groups. Subsequently, to obtain the spectra depicted in Fig. 1, we mixed these three ‘pure component’ spectra in accordance with the fitted ratio of the treated experimental 2DIR spectrum (i.e., a 0.6 : 0.21 : 0.19 *α*-helix : *β*-sheet : random-coil ratio).

## Results and discussion

### FTIR Spectrum

In Fig. 2 we show the infrared absorption spectra of untreated and hydrated silk films in the region between 1400 cm^*−*1^ and 1600 cm^*−*1^. In this region, the most predominant absorption bands are two vibrational bands of the amide group, amide I and amide II.^13^ The amide I mode originates mainly from the carbonyl stretching vibration, and absorbs between 1600 and 1700 cm^*−*1^, and the amide II is the ‘out-of-phase’ or asymmetric combination of C-N stretch and N-H bending, and absorbs between 1450 and 1550 cm^−1^. The untreated silk film shows a strong absorption band at 1650 cm^*−*1^, with a shoulder at lower frequency around 1630 cm^*−*1^. The presence of these bands indicates that silk contains more than one secondary structure. While the band at 1630 cm^*−*1^ can be assigned to *β*-sheet structures, ^13^ the assignment of the main band at 1650 cm^*−*1^ is not straightforward: both random-coil and *α*-helix structures can absorb at this frequency. ^13^ The amide II vibrational mode is found at lower frequency 1550 cm^*−*1^. In the region between 1400 cm^*−*1^ and 1500 cm^*−*1^, we also find different vibrational bands that we can assign to side-chain modes, such as CH bending. We later incubated the silk film in a saturated D_2_O environment for a period between 90 minutes at a RH of 85*±* 10%. The full measured infrared spectrum is also reported in Fig. S1. With increasing D_2_O exposure, the shoulder at 1630 cm^*−*1^ increases in intensity, and most of the amide II band at 1550 cm^*−*1^ red-shifts to around 1480 cm^*−*1^. We calculate the areas of the untreated and deuterated amide II bands, and find that *∼*70% of the NH groups were H/D exchanged to ND (see Fig. S2). Whereas it is clear that the exposure to a saturated D_2_O environment causes an increase of the shoulder at 1630 cm^*−*1^, suggesting, at first glance, an increase of the content of *β*-sheet structures. However, the congested infrared spectra do not directly provide information on the type of other secondary structures present in the film, and because of this, a quantitative estimation of the effect of increased humidity is not straightforward. To obtain a better understanding of the secondary structures in silk, we used two-dimensional infrared spectroscopy. Before discussing the 2DIR results, we briefly explain the principle of this spectroscopic method.

**Figure 2:**
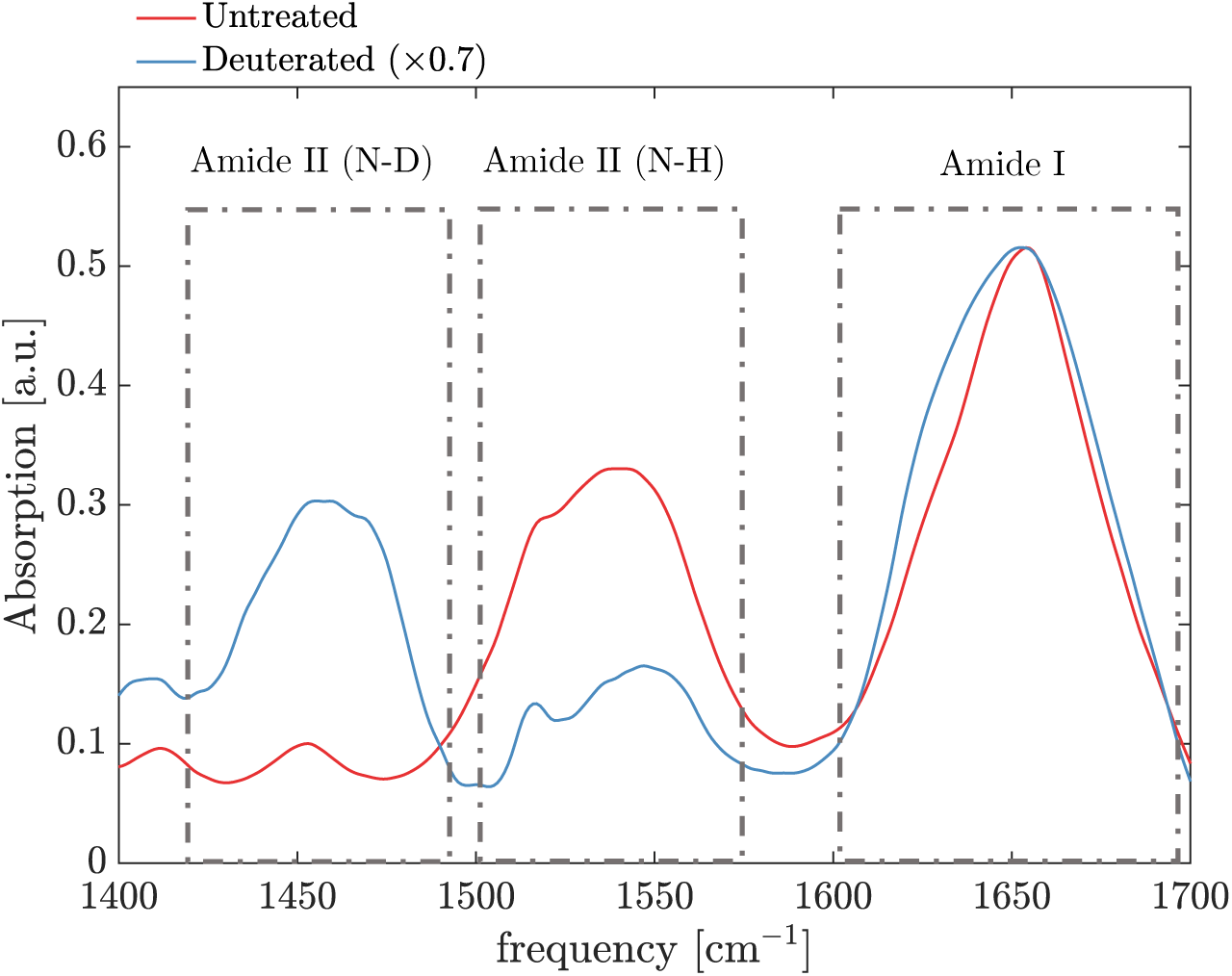
Infrared absorption spectra of a *∼*10 *µ*m thick silk film in the amide I and amide II absorption regions. The untreated sample was measured without further purification at ambient humidity (*∼*60 %). The untreated sample was then incubated at a RH (85 *±* 10%) in a saturated D_2_O environment for 120 minutes.

### Principle of 2DIR spectroscopy

In a 2DIR spectrum, vibrational couplings lead to specific spectral signatures (cross peaks) that contain information on the coupling strength (which depends strongly on the distance between the coupled vibrating bonds), and on the angle between the transition-dipole moments of the coupled modes.^23–25^ In pump-probe 2DIR spectroscopy, an intense, narrowband infrared pump pulse (with adjustable center frequency *ν*_pump_) resonantly excites at a specific frequency in the (in the present case, in the amide I band), and a delayed, broadband probe pulse is used to probe the resulting frequency-dependent IR-absorption change Δ*A*. Measuring the Δ*A* spectra for a range of *ν*_pump_ values, we obtain 2-dimensional spectra showing the pump-induced absorption change Δ*A*(*ν*_pump_, *ν*_probe_) as a function of the pump and probe frequencies *ν*_pump_ and *ν*_probe_. The 2DIR signal can be recorded with the pump- and probe-pulse polarizations parallel (||) or perpendicular (*⊥*) to each other, and the polarization-dependence of the cross-peak intensity is determined by the angle between the transition-dipole moments of the coupled modes.^25^

As an example, we show in Fig. 1c the simulated 2DIR spectra in the amide I region of a short peptide adopting ideal *β*-sheet and alpha-helix structure. When the pump-frequency is resonant with the *v* = 0 *→* 1 frequency of either of the two *β*-sheet modes (at *∼*1620 and 1670 cm^*−*1^), part of the molecules are excited to the *v* = 1 state of this mode, resulting in a decrease in the absorption at the *v* = 0 *→* 1 frequency (Δ*A <* 0 feature on the diagonal), and an increase in absorption at the *v* = 1 *→* 2 frequency (which is at a slightly lower value than the *v* = 0 *→* 1 frequency due to the anharmonicity of the vibrational potential), resulting in a Δ*A >* 0 feature slightly to the left of the diagonal. In this way, each normal mode gives rise to a +*/−* doublet on the diagonal of the 2D spectrum (diagonal peaks).

If two normal modes A and B are coupled, then (to first-order approximation) exciting mode A causes a small redshift of the frequency of mode B,^25^ resulting in an absorption decrease on the high-frequency side of the B band and an absorption increase on the low-frequency side, and so a +*/−* doublet at (*ν*_probe_, *ν*_pump_) = (*ν*_B_, *ν*_A_), of which the amplitude depends on the coupling strength. The intensities of the cross peaks also depends on the angle between the pump and probe polarizations, in a way that is determined purely by the angle between transition-dipole moments of the coupled modes. In particular, if the transition-dipole moments of the coupled modes are perpendicular (as is the case for the IR active modes of *α*-helices and *β*-sheets), then the relative intensity of the cross peaks is higher in the perpendicular spectrum than in the parallel spectrum (Fig. 1c). The set of diagonal and cross peak features (and its polarization dependence) of *α*-helices and *β*-sheets forms a pattern in the 2DIR spectrum that can be used as a finger print of these secondary structures.

### Polarization-dependent 2DIR spectra of *B. mori* silk

In Fig. 3a we show the perpendicular 2D-IR spectrum of the untreated silk sample. We observe strong diagonal peaks when exciting at 1660 cm^*−*1^ and 1630 cm^*−*1^. The diagonal peak at *v*_pump_ = 1630 cm^*−*1^ corresponds to the weak shoulder observed in the FTIR spectrum. In the 2DIR spectrum, this peak is better resolved because the 2DIR signal scales with square of the absorption cross-section *∼ σ*^2^, whereas the FTIR signal scales as *∼ σ*.^25^ The peaks colored in blue represent decreases in absorption (Δ*A <* 0) due to depletion of the amide-I *v* = 0 state, and the signal at lower probe frequency colored in red represents the induced absorption of the *ν* = 1 *→* 2 transition. In the off-diagonal region cross-peaks are visible. In particular, when exciting at 1630 cm^*−*1^, we observe a response at a probe frequency of 1700 cm^*−*1^. The negative half of the cross-peak +*/−* doublet is not clearly visible due to overlap with the positive part of the (much stronger) diagonal peak. Typically, the cross peaks in a 2DIR spectrum are much less intense than the diagonal peaks, and often they partly overlap with them. To isolate the cross peaks from the diagonal peaks we can use their different dependencies on the angle between the pump and probe polarizations. The diagonal peaks always have the same 3:1 intensity ratio for parallel vs perpendicular pump-probe polarizations, whereas for the cross peaks this intensity ratio is less than 3 (except in the special case where the transition-dipole moments of the coupled modes are exactly parallel).^25^ As a consequence, subtracting the perpendicular to parallel 2DIR spectra (with an appropriate scaling factor), we obtain a *diagonal free* 2DIR spectrum (see, as an example, the simulated spectrum in Fig. 1-(e)), and highlight only the cross peaks. ^39,40^ Furthermore, the magnitude of these cross-peaks is proportional to the amount of protein adopting a certain specific secondary structure. Thus, we can use the relative intensity of the crosspeak to monitor the relative change in secondary structure content. Removing the diagonal peaks by carefully subtracting perpendicular to parallel is especially useful for observing *α*-helix cross-peaks because these strongly overlap with the diagonal peaks (as the frequency difference between the two coupled modes A and E is only a few wavenumbers.^41,42^) In a similar fashion, provided that the parallel:perpendicular intensity ratio of all cross peaks in a 2DIR spectrum is the same, we can suppress the cross-peaks by subtracting the appropriately scaled perpendicular 2DIR spectrum from the parallel one, obtaining a *cross-peak free* 2DIR spectrum where only the diagonal peaks are present, as exemplified by the simulation in Fig. 1-(f).

**Figure 3:**
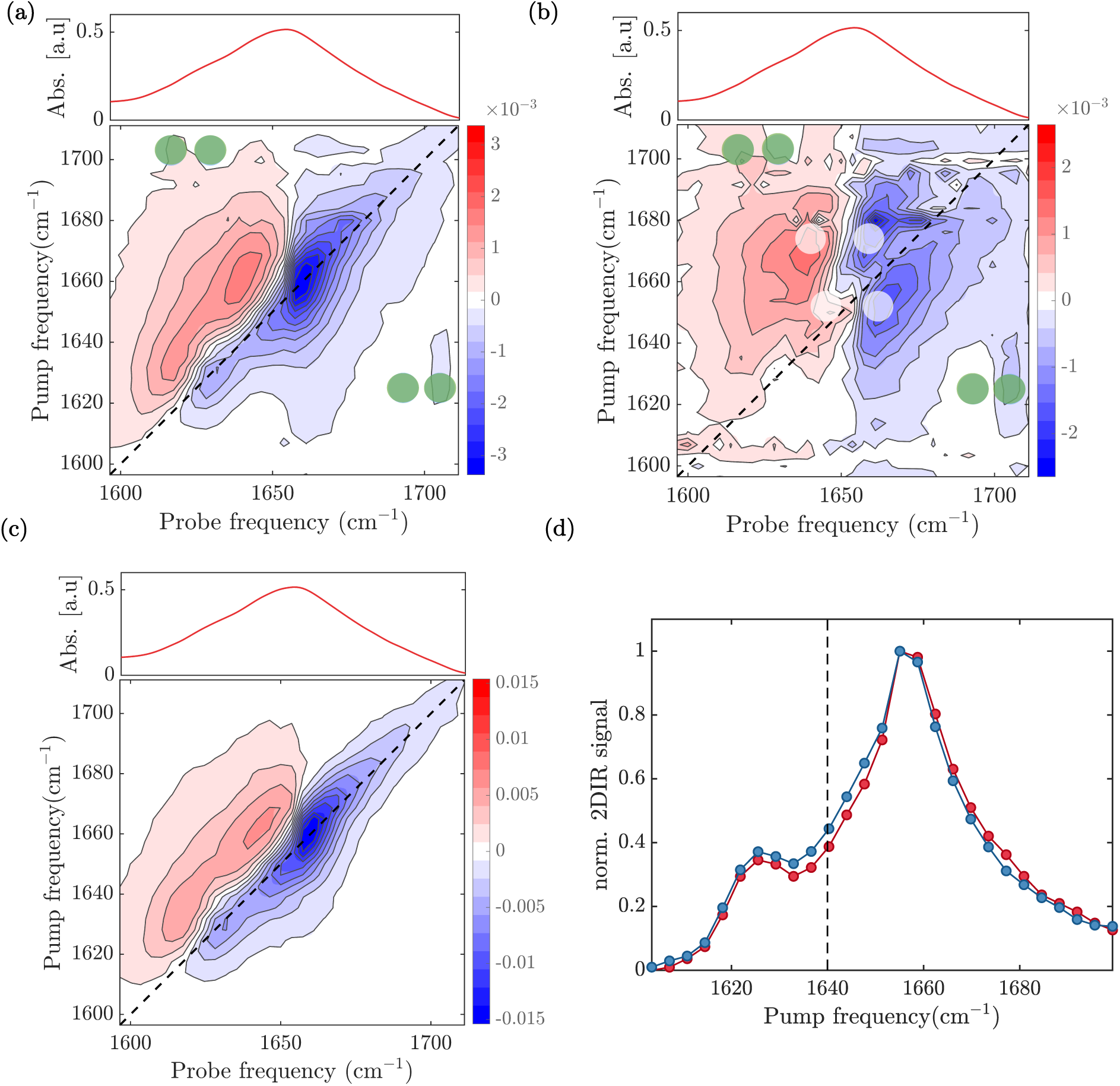
2DIR analysis of untreated silkworm film. a) Perpendicular and b) diagonal-free 2DIR spectra at a time delay of 1 ps. Green and white dots indicate the positions of *β*-sheet and *α*-helix cross-peaks, respectively. c) Cross-peaks free 2DIR spectrum at a time delay of 1 ps. d) Diagonal slices of the bleach signals of the isotropic (red circles) and cross-peaks free (blue circles) 2DIR spectra at a time delay of 1 ps. The absorption appears to be enhanced across quite a wide range of wavenumbers, from around 1630 to 1650 cm^*−*1^.

Fig.3-b shows the polarization-difference 2DIR spectrum obtained by subtracting parallel from 3 times perpendicular (in our experiments the parallel-to-perpendicular scaling factor is found to be 3 (Fig.S3), in agreement with the theoretical value). ^25^ We can now resolve the cross peak between 1630 and 1700 cm^*−*1^ (indicated by green dots), which indicate the presence of *β*-sheet structures. The 1630 cm^*−*1^ is the absorption band of the A_⊥_ mode, while the 1700 cm^*−*1^ is of the A_‖_ mode. Interestingly, we also observe cross-peak signatures when exciting at 1658 cm^*−*1^, and at 1665 cm^*−*1^ (indicated by the white dots). As previously explained, the coupling of the amide I modes in an *α*-helix leads to two delocalized modes, defined as A and E. In the linear infrared spectrum and in the isotropic 2DIR spectrum these two modes cannot be resolved, and give rise to a broad band centered around 1660 cm^*−*1^. However, since the A and E transition-dipole moments are perpendicularly oriented to each other, the visibility of the cross-peaks between the A and E modes is enhanced when subtracting the parallel signal from three times the perpendicular signal. This procedure reveals the presence of two cross peaks, with negative parts at *ν*_probe_ = 1658 cm^*−*1^ and at 1665 cm^*−*1^. The frequency splitting of the A and E modes is around 10 cm^*−*1^, in agreement with values reported in the literature. Fig. 3c shows the 2DIR spectrum obtained by subtracting perpendicular signal from 2 times parallel signal (to be compared with the simulated one in Fig. 1-f). As discussed before, by doing so, we eliminate the cross-peaks between perpendicular modes. In Fig. 3-d, we show the diagonal slices of the bleach signals of the isotropic and cross-peak free 2DIR spectra. The cross-peak-free diagonal slice reveals the present of a third band around 1650 cm^*−*1^, which we assign to random-coil structure. Fig. 4a-b shows the perpendicular and polarization-difference 2DIR spectra of hydrated silk film, respectively. We again observe the presence of two main diagonal peaks, obtained when exciting at 1630 and 1660 cm^*−*1^. The peaks are much more elongated because of the increased inhomogeneous broadening, reflecting the structural disorder caused by the increased hydration of the amide groups.^25^ In the diagonal-free 2DIR spectrum Fig. 4-b, we notice the same cross-peaks features observed previously, indicating that upon hydration the silk film maintains *α*-helical and *β*-sheet conformations. We observe, however, that the increased hydration lowers the frequency of the highest vibrational band and broadens the cross-peak of the *β*-sheet structure: in the untreated sample, the A_‖_ absorbs at 1705 cm^*−*1^, while in the hydrated sample at 1700 cm^*−*1^. The frequency shift and broadening indicate that part of the *β*-sheet structures become hydrated, likely in the interphase part between crystalline and amourphous regions. ^20^ To better resolve the diagonal signatures, we again subtract scaled parallel to perpendicular to remove the cross-peaks (Fig. 4-b). The diagonal slices of the cross-peak free and isotropic signal again reveal the presence of three bands (1630 cm^*−*1^, 1640 cm^*−*1^ and 1660 cm^*−*1^).

**Figure 4:**
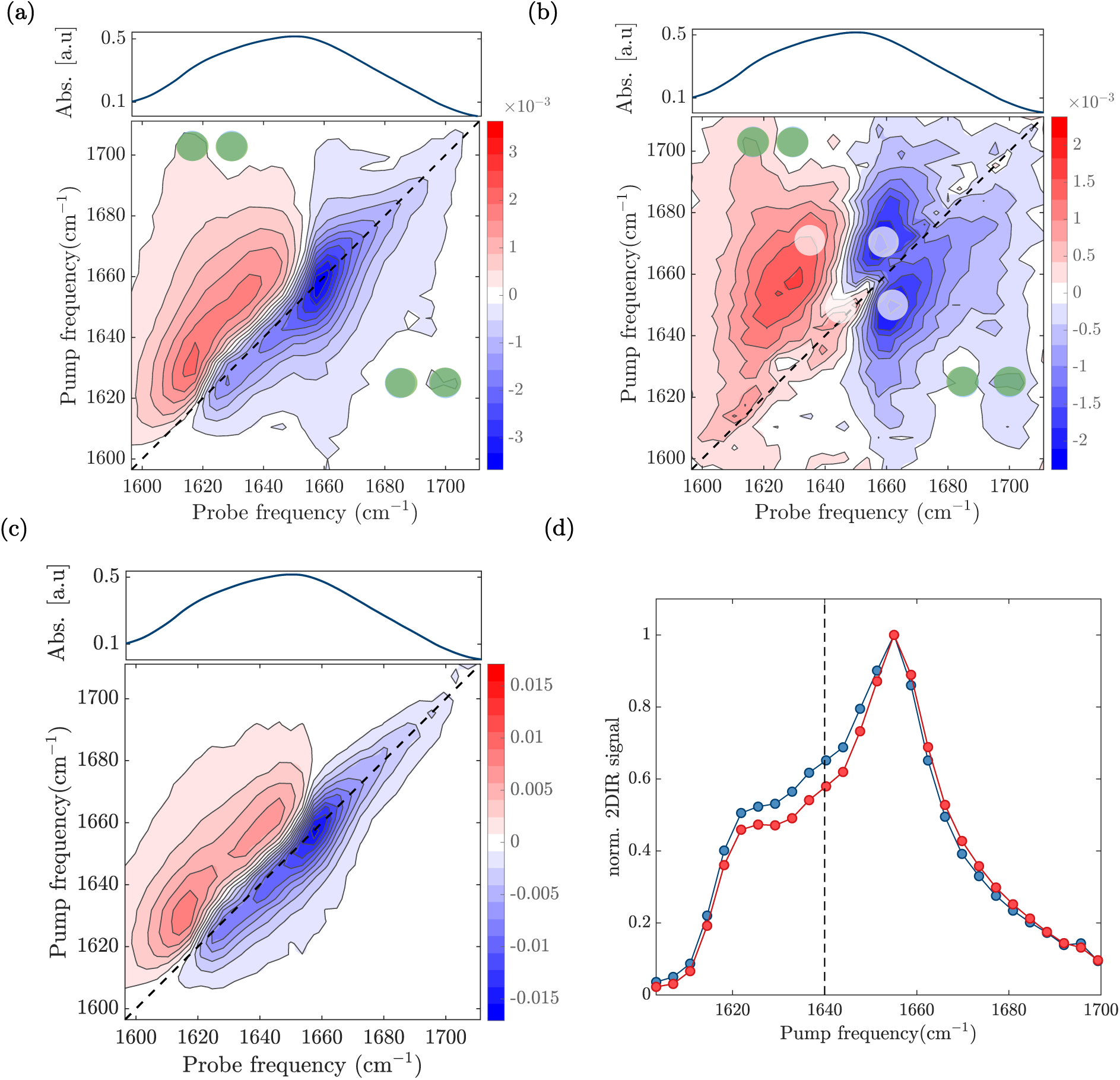
2DIR analysis of treated silkworm film. a) Perpendicular and b) diagonal-free 2DIR spectra at a time delay of 1 ps. Green and white dots indicate the positions of *β*-sheet and *α*-helix cross-peaks, respectively. c) Cross-peaks free 2DIR spectrum at a time delay of 1 ps. The linear absorption spectra of the treated silkworm film is reported on top of the 2D-IR spectra (a,b,c) for comparison. d) Diagonal slices of the bleach signals of the isotropic (red circles) and cross-peaks free (blue circles) 2DIR spectra at a time delay of 1 ps. Vertical line shows the enhanced absorption from 1620 till 1650 cm^*−*1^.

In Fig.5-(a), we report the cross-peak free diagonal slices of the treated and untreated samples, which we normalize to the respective spectrum areas. Compared to the linear infrared spectra, we can now better resolve the individual vibrational bands. This is because the 2DIR signal scales as the cross-section *σ*^2^, while the linear IR signal scales as *σ*, leading to narrower peaks.^43^ In the untreated spectrum, the 2DIR signal shows a strong absorption band centered at 1660 cm^-1^, which we assigned before to *α*-helix structure. At ambient humidity, we find that the dominant secondary structure is *α*-helix (this is reproduced in a second sample, see Fig. S4). Upon increasing the humidity, the *α*-helix peak decreases in intensity while the 2DIR signal around 1640 and 1620 cm^-1^ increases. The enhancement of the absorption at this lower frequency region may be due to an increase in *β*-sheet and/or random coil contents. In this case, determining the relative contribution of these two secondary structures to such increase is not easy. In fact, the two vibrational bands strongly overlap with each other, and there is no major change at one single absorption frequency, specific either for the random coil or *β*-sheet mode. Multipeak fitting analysis will, hence, lead to ambiguous results in this case. To solve this problem, we use the diagonal-free 2DIR spectra, where the *β*-sheet cross-peaks are well-resolved. By studying their relative intensity, we can directly estimate whether the *β*-sheet content increases since the magnitude of these crosspeaks scales with the *β*-sheet content. Fig.5-(b) shows the anti-diagonal slices obtained from the diagonal free 2DIR spectra (Fig. S5). We normalize them to the area of the respective cross-peak free diagonal slices. We observe that both *β*-sheet cross-peaks absorbing around 1625 and 1700 cm^-1^ do not change significantly, suggesting that the *β*-sheet content stays constant, whereas the random-coil increases. Longer exposure to humidity leads to a clear increase of the *β*-sheet structure (see Fig. S6), in agreement with previous studies. ^44^ These previous studies suggest that water weakens the hydrogen bonds within the *α*-helix, enabling chain movement and *β*-sheet formation because of helix-helix interactions, similar to what happens in amyloid formation. The fact that we observe an increase of random-coil at short exposure time to D_2_O might suggest the possibility that the conversion *α*-helix to *β*-sheet structure requires a critical hydration level, which it is not reached in the first 2 hours. However, the hydration might be enough to lead to the unfolding of the *α*-helix into random coil. ^44^

**Figure 5:**
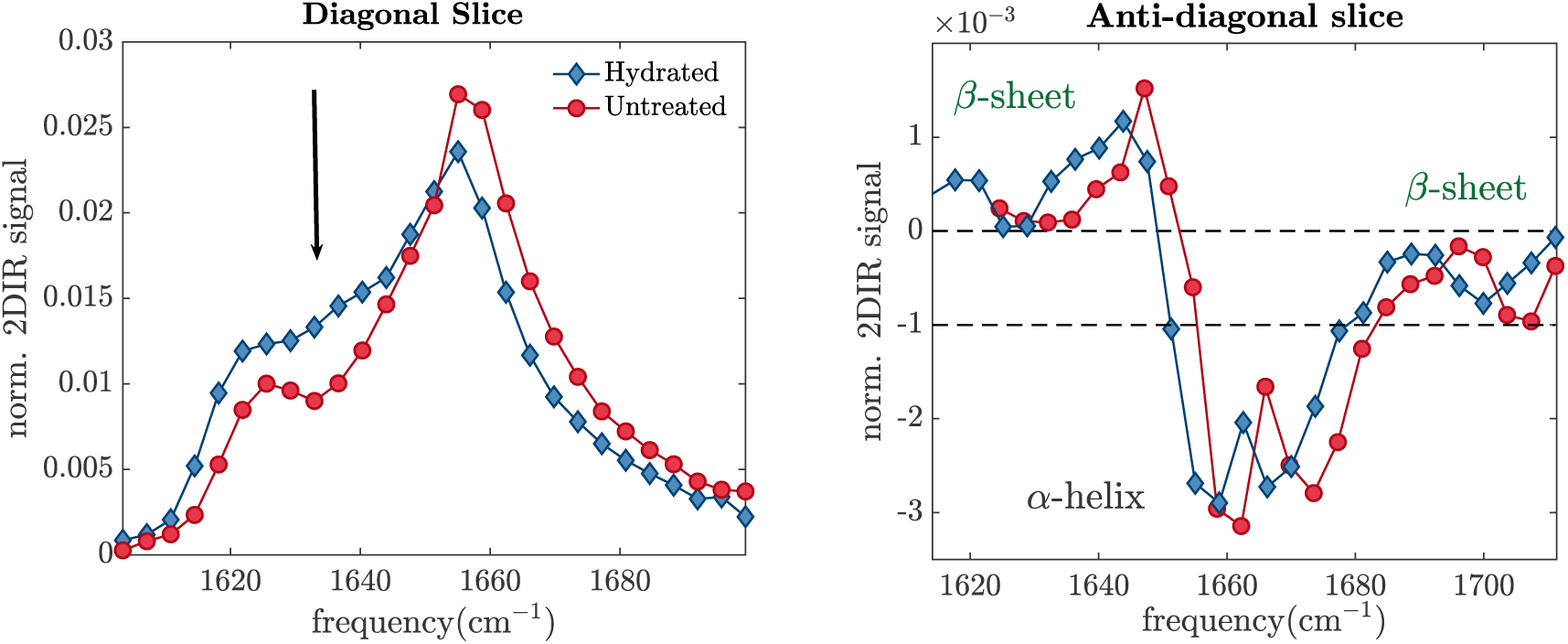
Untreated and treated isotropic diagonal (a) and anti-diagonal slices (b). (a): diagonal slices obtained from the 2D-IR spectra shown in Fig. 3-(c) and Fig. 4-(c), normalized in area. (b): Anti-diagonal slices of the 2D-IR spectra shown in Fig. 3-(b) and Fig. 4-(b). The spectra have been normalized to the area of the diagonal-free spectrum.

In light of our results, we caution researchers using multi-peak fitting analysis into making the assumption, based on previous studies, that an increase in the vibrational region between 1620-1640 cm^-1^ is mostly determined by the increase of *β*-sheet content, especially at the early stages of the hydration process.

## Conclusion

In this paper, we show that 2DIR spectroscopy can be used to disentangle the secondary structures in a complex and unpurified bio-material such as silk film in a label-free and not invasive manner. From polarization-difference 2DIR spectra with the appropriate weighting factors, we obtain 2DIR spectra where we identify and isolate or cross-peak or diagonal peaks. Because *α*-helix and *β*-sheet structures have specific cross-peak patterns, we can assign vibrational bands to specific secondary structures. In the same way, by exploiting the same polarization dependency, we can also remove the cross-peaks that overlap with diagonal peaks, reducing the spectral congestion in 2DIR, and thus enhancing our spectral resolution. This enables us to resolve the presence of an additional vibrational band, which it is assigned to random-coil. Thanks to this enhanced resolution, we find that at ambient humidity the dominant conformation is *α*-helix in the film studied here, while *β*-sheet and random coil structures are present with lower abundance. Upon exposure to high humidity, we find that *α*-helix content decreases, while the content of *β*-sheet/random-coil increases. By comparing the relative magnitudes of the *β*-sheet cross-peaks, we find that the *β*-sheet content does not significantly change, implying that, in the sample analyzed in this paper, *α*-helix mainly converts to random coil. We thus show that (1) we can disentangle the 2DIR spectra of unpurified silkworm films, resolving the presence of definite vibrational bands, and, in case of *β*-sheet and *α*-helix,(2) assign the vibrational bands to specific secondary structures and (3) determine their relative change when exposing silkworm film to humidity without further data analysis.

The molecular properties of the building blocks of hierarchical biomaterials determine the physical properties that are required to fulfill their biological functionalities. Understanding the connection between molecular and macroscopic properties is thus a key determinant to elucidate the success and failure of biomaterials. Here, we showed that 2DIR can be applied successfully to provide novel, unique and specific structural information about the building blocks of unpurified biomaterials. Further application of 2DIR in combination with other techniques, such as rheology, should enable us to gain a better understanding of the critical relationship between biomechanical and biomolecular properties.

## Acknowledgement

We thank Prof. Dr. Gijsje Koenderink for helping us initiate the collaboration that gave rise to this article and Prof. Dr. Daniel Bonn for the fruitful discussions. F.C. acknowledges financial support from The Netherlands Organization for Scientific Research (NWO) (Grant Number 680-91-13). A.S.C and T.W acknowledge support by the Novo Nordisk Foundation (Facility Grant NanoScat, no. NNF18OC0032628)

## Supporting Information Available

Simulated parallel and isotropic 2DIR spectrum; Linear IR spectrum; Amide II spectral fit; Comparison 2DIR signal/IR signal

