## Supplementary text and figures for "In situ identification of secondary structures in unpurified *Bombyx mori* silk fibrils using polarized two-dimensional infrared spectroscopy"

### Supplementary for: In situ identification of secondary structures in unpurified silkworm films using polarized advanced infrared spectroscopy

Giulia Giubertoni,[*,†,‡,‖] Federico Caporaletti,[*,‡,†,‖] Steven Roeters,[¶,†] Adam S. Chatterley,[¶] Tobias Weidner,[¶] Peter Laity,[§] Chris Holland,[§] and Sander Woutersen[†]

† *Van 't Hoff Institute for Molecular Sciences, University of Amsterdam, Science Park 904, 1098XH Amsterdam, The Netherlands*

‡ *Van der Waals-Zeeman Institute, Institute of Physics, University of Amsterdam, 1098XH Amsterdam, The Netherlands*

¶ *Department of Chemistry, Aarhus University, Aarhus C, Denmark*

§ *Department of Materials Science and Engineering, University of Sheffield, Sir Robert Hadfield Building, Mappin St., Sheffield S1 3JD, UK*

‖ *These authors contributed equally to the work*

### FTIR spectra of the hydrated silkworm films

Fig. S1 shows the extended IR-spectra of the untreated and hydrated silkworm films. The silkworm films have been exposed to an environment saturated with $D_2O$ instead of $H_2O$ to suppress the spectral contributions of the H-O-H bending mode of water, which would have masked the amide region of the spectrum. After the treatment a broad vibrational band, correspnding to the O-D stretching of water, appears around 2500 cm$^{-1}$. The amide groups

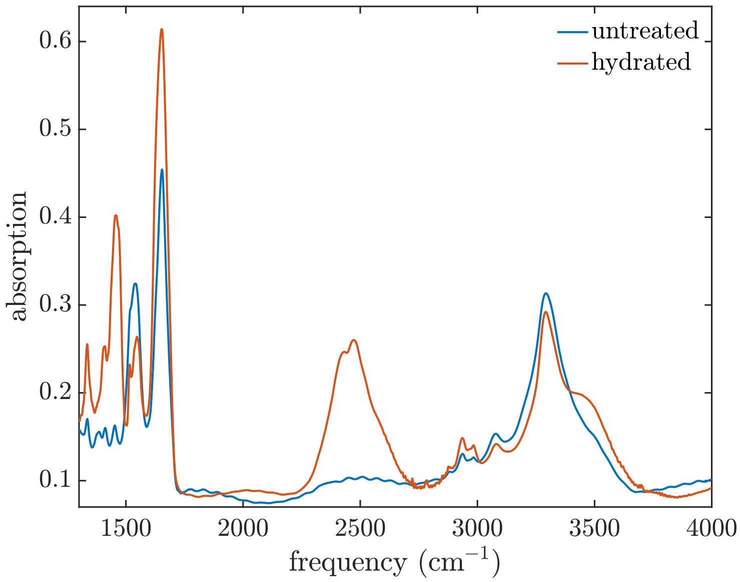


Figure S1: Linear infrared spectra hydrated and untreated silk film produced by Bombyx Caterpillars. The exposure to a environment saturated of $D_2O$ leads to the appearance of the O-D stretch vibrational band around 2500 cm$^{-1}$.

that are able to isotopically exhange H with D during the hydration process are those that are water accessible. The area of the vibrational band of the N-D amide II hence reflects the amount of groups in the film that can interact with water. Fig. S2 shows the N-D and N-H amide II vibrational bands for Sample A discussed in the main text: from the calculated areas we can estimate that $\simeq 70\%$ of the NH groups are H/D exchanged to ND.

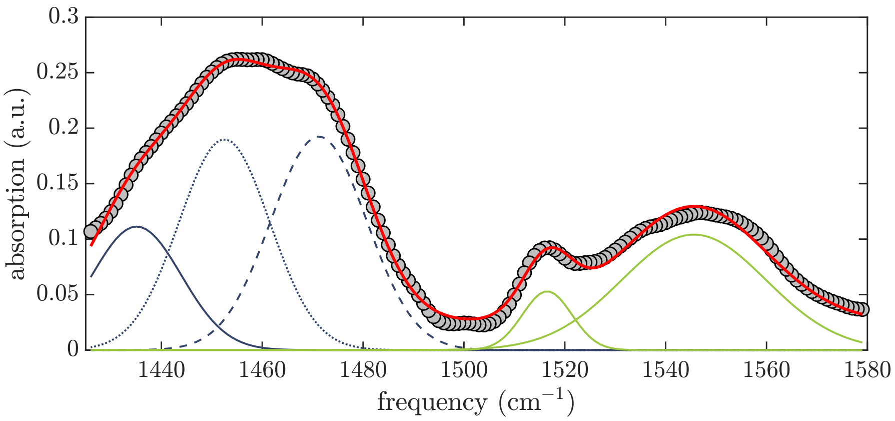


Figure S2: Fit of the linear spectrum of hydrated silk film in the amide II region. To estimate the amount of amide groups that isotopically exchange (i.e. that are water accessible), we extract the areas of vibrational bands of the N-D amide II, and N-H amide II. To obtain a quantitatively good fit, we use three Gaussian-shaped peaks to describe the amide II absorption bands at 1450 cm$^{-1}$, and 2 Gaussian-shaped peaks to described the amide II at 1550 cm$^{-1}$.

### Anisotropy

To obtain information over the molecular orientation, we calculate the anisotropy, defined as $R = \frac{\Delta\alpha_{\text{par}} - \Delta\alpha_{\text{per}}}{\Delta\alpha_{\text{par}} + 2\Delta\alpha_{\text{per}}}$, where $\Delta\alpha_{\text{par}}$ and $\Delta\alpha_{\text{per}}$ are the transient absorption changes measured in parallel and in perpendicular polarization configuration, respectively. In case of the diagonal peaks, the ratio between parallel and perpendicular signal is expected to be 3, leading to an anisotropy of 0.4. In Fig.S3, we report the anisotropy as a function of probe frequency obtained by centering the excitation pulse at the beta-sheet $A_\perp$ vibrational mode, which absorbs at 1623 cm$^{-1}$. We observe that at the probe frequency of the same mode, where the bleach of the diagonal peak is found in the 2DIR spectrum, the anisotropy value is perfectly at 0.4, indicating that the scaling factor between parallel and perpendicular is 3 in our experiments. We also observe that at 1710 cm$^{-1}$, where the $\beta$-sheet $A_\parallel$ vibrational mode absorbs, the anisotropy changes to negative values, approaching -0.2. In this case, the anisotropy represents the relative orientation of the $A_\perp$ transition dipole moment with respect to the $A_\parallel$ dipole moment. We can calculate the relative angle $\theta = \arccos\sqrt{\frac{5R_0+1}{3}}$, obtaining an angle of around 90°, which is expected since $A_\perp$ and $A_\parallel$ have transition dipole moments that lie perpendicular to each other.

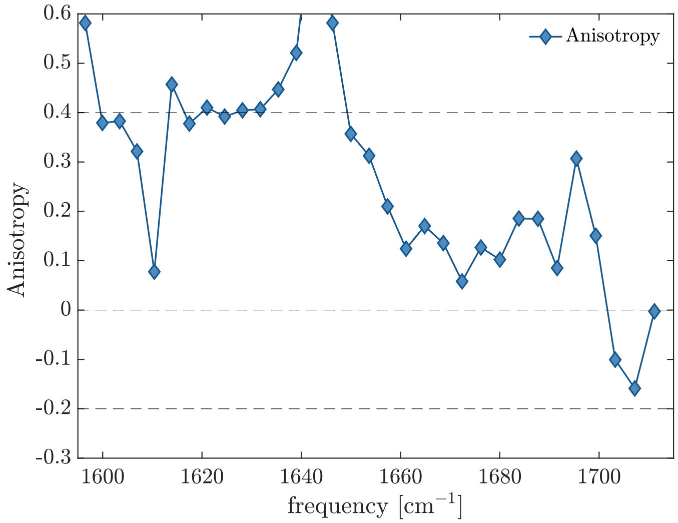


Figure S3: Anisotropy values as a function of probe frequency upon excitation of th e $\beta$-sheet $A_{\perp}$ vibrational mode. At certain frequencies, such as 1645 cm$^{-1}$, the anisotropy values are out of scale because these are the frequencies of the nodal line between the negative and positive peaks, where the parallel and perpendicular signals are $\sim 0$.

### Absence of $\beta$-sheet secondary structures in untreated films

Fig. S4 shows the "diagonal-free" spectrum of a sample taken from another silk film before being hydrated. We here observe only the off-diagonal signatures of the $\alpha$-helix, confirming that in silkworm films proteins adopt mostly $\alpha$-helix structure at ambient conditions.

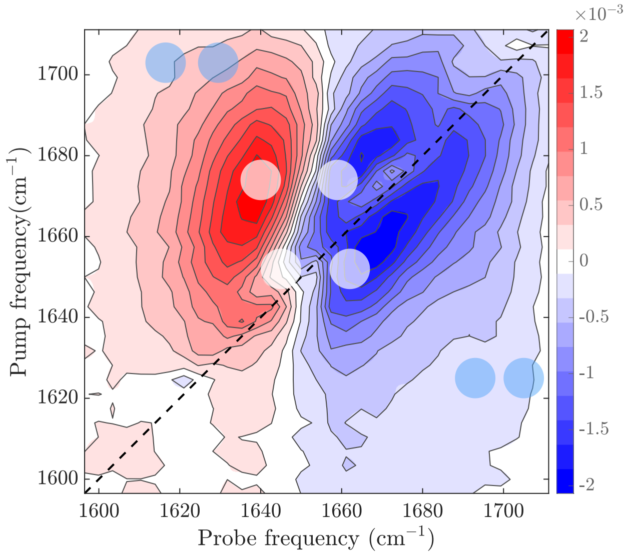


Figure S4: Subtracted 2D-IR spectrum of an untreated silk film. The spectrum displays the cross-peaks associated to $\alpha$-helix secondary structures (see white circles), while the cross-peaks signatures expected for $\beta$-sheet, which would be expected at the frequencies highlighted by the blue circles, are absent.

### Anti-diagonal slices

In Fig.S5 we report the diagonal free 2DIR spectra and the anti-diagonal slices from where we obtain the 2DIR signals reported in the main text.

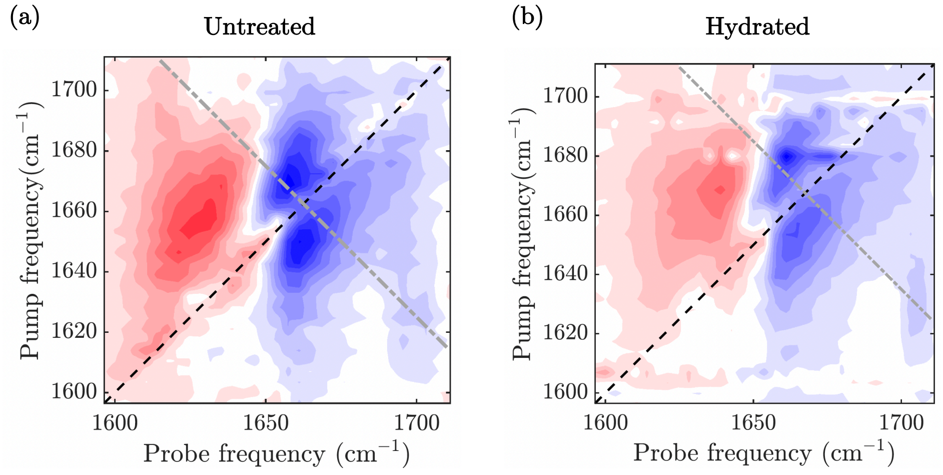


Figure S5: Diagonal free 2D-IR spectra of the untreated (a) and hydrated (b) films discussed in the main text. The main diagonal (black dashed line) and anti-diagonal slice (gray dashed-dotted line), used in Fig. 5 are highlighted.

### Effect of exposure time to high humidity on $\beta$-sheet content

Data reported here were taken on the at the University of Aarhus using a 10 kHz commercial time-domain 2DIR spectrometer (PhaseTech 2DQuickIR) described previously.[1,2] Briefly, femtosecond mid-IR pulses were split into pump and probe pulses. The pump pulses were split into time- and phase-controlled pulse pairs using an acousto-optic pulse shaper, and then focused into the sample 500 fs before the probe pulse, which was dispersed onto an MCT detector (PhaseTech JackHammer). The delay between the pump pulse varied in 33 fs steps from 0 to 3 ps, and the measurements were performed in a 1300 cm-1 rotating frame. In order to reduce interference from scattered light, a 4-frame phase cycling scheme was used, and reference spectra recorded at -10 ps were subtracted. The sample here analyzed is taken from the same silkworm film on which the experiments reported in the main text were performed. After exposing the film to saturated $D_2O$ (85% RH) no major change is observed in magnitude of the cross-peaks between the $\beta$-sheet modes at 1630 cm$^{-1}$, and at 1700 cm$^{-1}$, indicating that the $\beta$-sheet content is constant. This confirms and well-reproduces the result presented in the main text. We further expose this sample to high humidity for additional

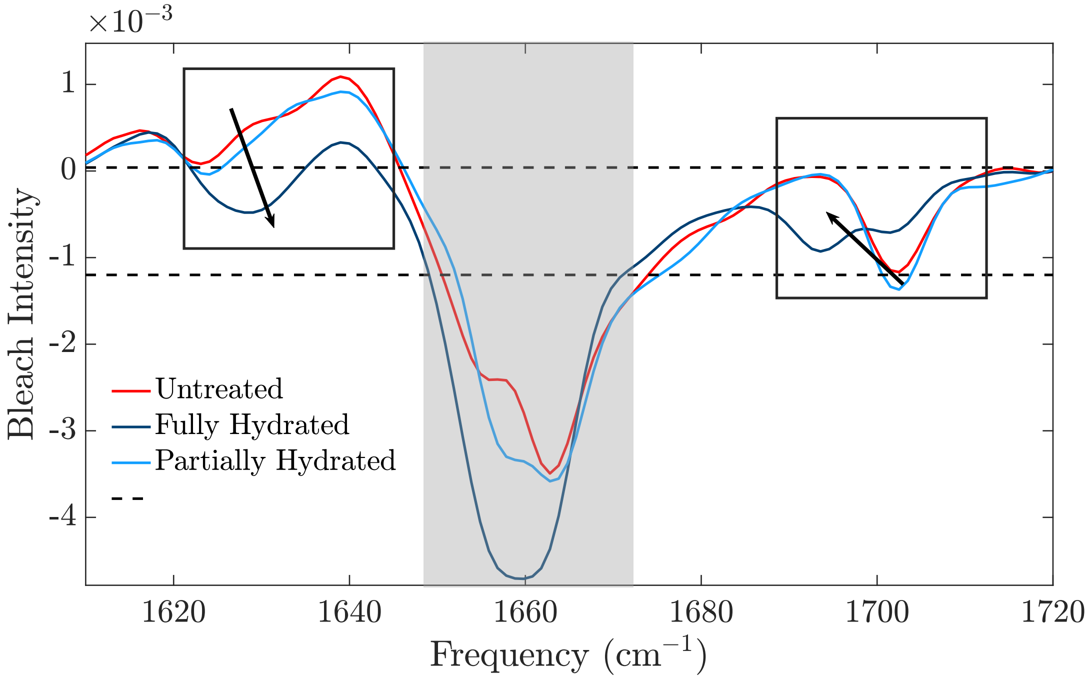


Figure S6: Anti-diagonal slice of the diagonal free spectrum measured from the same silkworm film as in the main text at different level of hydration: untreated (red solid line), hydrated for 2 hours (cyan solid line) and for 36 hours (blue solid line). The black boxes highlights the changes occurring in the cross-peaks associated to $\beta$-sheet secondary structures. The gray rectangle shadows the $\alpha$-helix region, close to the main diagonal, that is affected by scattering in the shown measurement.

34 hours. Upon this, the cross-peaks between the $\beta$-sheet modes change: the magnitude of the cross-peak at 1620 cm$^{-1}$ increases drastically, while the cross-peak at 1700 cm$^{-1}$ spits in two distinct subbands. This last effect is due to the overlap of different cross-peaks, likely associated to the hydrated and not-hydrated $\beta$-sheet. Because of the overlap, the relative intensities of the cross-peaks cancel partially off, and thus we do not consider it in our analysis. However, it is clearly visible that the magnitude of the cross-peak at 1620 cm$^{-1}$ increases drastically, indicating that $\beta$-sheet structures are being formed after long exposure time (see blue curve in Fig. S6).
